# Catching up to a fast-moving target: Evaluation of a health system strengthening intervention in rural Rwanda 2005-2010 using data from repeated cross-sectional surveys

**DOI:** 10.1101/238543

**Authors:** Dana R. Thomson, Cheryl Amoroso, Sidney Atwood, Matthew H. Bonds, Felix Cyamatare Rwabukwisi, Peter Drobac, Karen E. Finnegan, Didi Bertrand Farmer, Paul E. Farmer, Antoinette Habinshuti, Lisa R. Hirschhorn, Anatole Manzi, Peter Niyigena, Michael L. Rich, Sara Stulac, Megan Murray, Agnes Binagwaho

**Author notes:** Joint senior authors (Megan Murray, Agnes Binagwaho).

## Abstract

**Introduction:** Although Rwanda’s health system underwent major reforms and improvements after the 1994 Genocide, the health system and population health in the southeast lagged behind other areas. In 2005 Partners In Health and the Rwandan Ministry of Health began a health system strengthening intervention in this region.

**Methods:** Combining results from the 2005 and 2010 Demographic and Health Surveys with those from a supplemental 2010 survey, we compared changes in health system output indicators and population health outcomes between 2005 and 2010 as reported by 21,338 women living in the intervention area and similar rural areas, controlling for potential confounding by economic and demographic variables.

**Results:** Overall health system coverage improved similarly in both regions between 2005 and 2010, with an indicator of composite coverage of child health interventions increasing from 57.9% to 75.0% in the intervention area and from 58.7% to 73.8% in other rural areas. Despite experiencing poorer health outcomes in 2005, the intervention area caught up to or exceeded other rural areas on 23 of 25 indicators. Most notably, under-five mortality declined by an annual rate of 12.8% in the intervention area, from 229.8 to 83.2 deaths per 1000 live births, and by 8.9% in other rural areas, from 157.7 to 75.8 deaths per 1000 live births. Improvements were most dramatic among the poorest households.

**Conclusion:** We observed dramatic improvements in population health outcomes including under-five mortality between 2005 and 2010 in rural Rwanda generally, and in the intervention area specifically.

**SUMMARY BOX:** *What is already known about this topic?:*

- Much of the evidence that health system strengthening in rural Africa has improved health outcomes comes from studies of targeted regional interventions such as performance based financing or community health worker programs, rather than integrated interventions that encompass multiple components including infrastructure and supply chain investments, health management information system, workforce training and incentives at all levels, community health workers, and free services for poor patients.
- In addition to these experimental or quasi-experimental studies, a series of case studies have documented individual nations, pathways to achieving millennium development goal 4 target, the reduction of under-five mortality by two thirds between 1990 and 2015.
- These reports suggest that improvements in coverage of reproductive, maternal and child health indicators explain some, but not all, of the decline in child mortality and that these successes occurred in the context of national gains in health, nutrition and food security, sanitation, poverty reduction, and access to clean water.

*What are the new findings?:*

- Coverage of most maternal and child health care interventions improved at a similar pace in our rural intervention area and other rural areas.
- Despite experiencing poorer health outcomes in 2005, our rural intervention area caught up to or exceeded other rural areas on 23 of 25 population health indicators by 2010.
- Infant and under-5 mortality declined in our rural intervention area even more precipitously than in other rural areas of Rwanda between 2005 and 2010.

*How might this influence practice?:*

- The process of strengthening national health systems often involves trade-offs between a focus on first testing individual programs that distributed widely, as is often practiced by pilot programs with multilateral institutions, or implementing multiple simultaneous programs locally. Our results show that integrated health system strengthening interventions can be locally adapted to enable the rapid expansion of health care coverage as well as dramatic improvements in population health outcomes.
- Integrated multi-level interventions can also help narrow the health care coverage and outcome gap between richer and poorer members of a society.
- National governments can leverage nongovernmental partners to achieve the health related sustainable development goals through joint implementation of national health policy.

## INTRODUCTION

The 1994 Rwanda Genocide was followed by a profound decline in population health that persisted for almost a decade. In the aftermath of the killing of nearly 20% of the population, HIV incidence soared, a cholera epidemic among the Rwandan refugees ensued, and vaccination rates plummeted. In 2000, the new government launched a development initiative, Vision 2020, of which health equity was a major component, and in 2003, Rwanda established health as an inalienable right.^1^ The many health care initiatives implemented nationally between 2003 and 2010 included a national health insurance policy,^2^ performance based financing of health programs,^3^ a village community health worker program,^3^ scale up of vaccinations,^4^ HIV treatment,^5^ and malaria reduction initiatives.^6^ Between 2004 and 2011, ART coverage increased seven fold to 94.0% and by 2011,^7^ the government of Rwanda (GoR) spent 10% of public expenditure on health.^8^

In 2005, the non-governmental organization, Partners In Health (PIH), and the Rwandan Ministry of Health (RMOH) began a collaboration to strengthen the health system in a region of southeastern Rwanda (henceforth referred to as Kirehe/S. Kayonza) where health outcomes were among the worst in Rwanda. Children in this area experienced higher rates of death, acute respiratory infection (ARI), diarrhea, and fever than in all the other rural areas of the country combined.^9^ To address this, PIH and RMOH jointly led a regional effort based on the World Health Organization (WHO) six building blocks of health system strengthening.^10^ The intervention included the renovation and equipping of dysfunctional health facilities, the recruitment, retention, and training of a health workforce, the development of a medical record system, the procurement of medical products and technologies, financial support to offset health insurance premium costs and user fees, and the development of governance strategies to ensure the longevity of the project. These interventions coincided with major RMOH reforms to coordinate external aid with government policies, scale-up a community-based health insurance scheme, and introduce performance-based pay into the district health system.^11^ Specific aspects of the PIH-RMOH intervention are described in the Supplement and reviewed elsewhere.^12^

An important principle of the RMOH-PIH collaboration in Kirehe/S. Kayonza was to leverage existing resources rather than spend limited resources on building new systems. Routinely conducted Demographic and Health Surveys (DHSs) provide extensive data that can be used to measure health system outputs and population health outcomes.^13^ We evaluated the impact of the RMOH-PIH interventions by comparing the temporal trends in health outputs and outcomes between 2005 and 2010 in the target region to those in other rural areas.

## METHODS

### Indicators

We assessed the following health system *output* indicators: whether treatment was provided for recent episodes of acute respiratory infection (ARI), diarrhea, or fever in children under age five; whether children under age two received the recommended three doses of DPT or measles vaccine; whether children received vitamin A supplementation between age six months and one year; whether children were exclusively breastfed for the first six months of life; whether at least one, or the recommended four, antenatal care visits took place during the last pregnancy; whether the most recent birth was attended by a skilled health worker; whether the birth was delivered by Cesarean-section; whether women received postnatal care within 24 hours of delivery; women’s current contraceptive use; and their unmet need for contraception, and the following population health *outcome* indicators: neonatal, infant, and under-five year old mortality; adult mortality (men, women, and combined); recent occurrence of ARI, diarrhea, or fever in children under age five; and stunting and wasting in children under age five. ^14^

### Data

We used Rwanda Demographic and Health Survey (RDHS) data collected from 21,338 women living in Kirehe/S. Kayonza (K/SK) and other rural areas (ORA) in 2005 (K/SK: 418, ORA: 8,217) and 2010 (K/SK: 2,073, ORA: 10,630) (Table 1). RDHSs are nationally and sub-nationally representative two-stage cluster samples conducted roughly every five years by the RMOH, National Institute of Statistics-Rwanda (NISR), and ICF International. The surveys collect information from women aged 15 to 49 on their reproductive health histories, practices, and desires; household composition; siblings’, survival; and children’s health and survival. The DHS birth history module included the date of the birth and death of each child born alive, and through the sibling module, the age and date of death for each biological sibling (see Supplement). The 2005 RDHS was underway at the onset of the RMOH-PIH collaboration. We coordinated with the NISR immediately following the 2010 RDHS to collect a supplemental sample of 1,391 households from 54 primary sampling units (PSUs) in Kirehe/S. Kayonza using the same sampling frame, staff, and questionnaires as the 2010 RDHS (Figure 1).^12^ Most data were collected consistently across the three surveys (2005 RDHS, 2010 RDHS, Supplemental RDHS) although neither stunting and wasting were assessed in the supplemental survey.

**Table 1.** Summary of socio-demographic characteristics in 2005 and 2010

|  | **2005** |  | **2010** |  | **Comparison of group trends** |
| --- | --- | --- | --- | --- | --- |
|  | ORA | K/SK | ORA | K/SK | p-value‡ |
| **Women (15-49)** |  |  |  |  |  |
| Age | 28.6 | 28.1 | 28.6 | 28.7 | 0.059 |
| Married/partnered | 50.6% | 54.1% | 51.2% | 54.4% | 0.904 |
| Employed | 94.6% | 94.6% | 86.1% | 87.9% | 0.379 |
| Literacy | 68.0% | 63.7% | 75.2% | 73.3% | 0.454 |
| **N (weighted)** | 8,877 | 523 | 10,957 | 2,084 |  |
| **N (unweighted)** | 8,217 | 418 | 10,630 | 2,073 |  |
| **Households** |  |  |  |  |  |
| Wealth score† | -0.246 | -0.290 | 0.242 | 0.293 | 0.013 |
| Electricity | 1.4% | <0.1% | 4.1% | 4.5% | 0.175 |
| Finished floor | 7.1% | 5.1% | 10.7% | 8.5% | 0.902 |
| Improved water | 27.9% | 47.5% | 72.4% | 65.5% | 0.014 |
| Members 15+ with primary education | 66.7% | 70.5% | 75.2% | 75.2% | 0.205 |
| **N (weighted)** | 8,168 | 498 | 10,157 | 2,041 |  |
| **N (unweighted)** | 7,702 | 394 | 9,891 | 2,031 |  |
| **Primary Sampling Units** |  |  |  |  |  |
| Distance to road (meters) | 3105 | 3275 | 3221 | 2960 | 0.543 |
| Distance to Kigali (km) | 50.8 | 49.6 | 47.2 | 46.3 | 0.972 |
| Elevation (meters) | 1763 | 1488 | 1800 | 1527 | 0.990 |
| Jan average rainfall (mm) | 113 | 117 | 113 | 115 | 0.483 |
| Apr average rainfall (mm) | 175 | 176 | 175 | 176 | 0.965 |
| Jul average rainfall (mm) | 83 | 67 | 83 | 69 | 0.507 |
| Oct average rainfall (mm) | 188 | 179 | 188 | 180 | 0.417 |
| **N (unweighted)** | 331 | 17 | 388 | 79 |  |

† With respect to 2005 wealth score definition
‡ Based on t-test from an OLS regression with year, group, and year-group interaction terms

**Figure 1.**
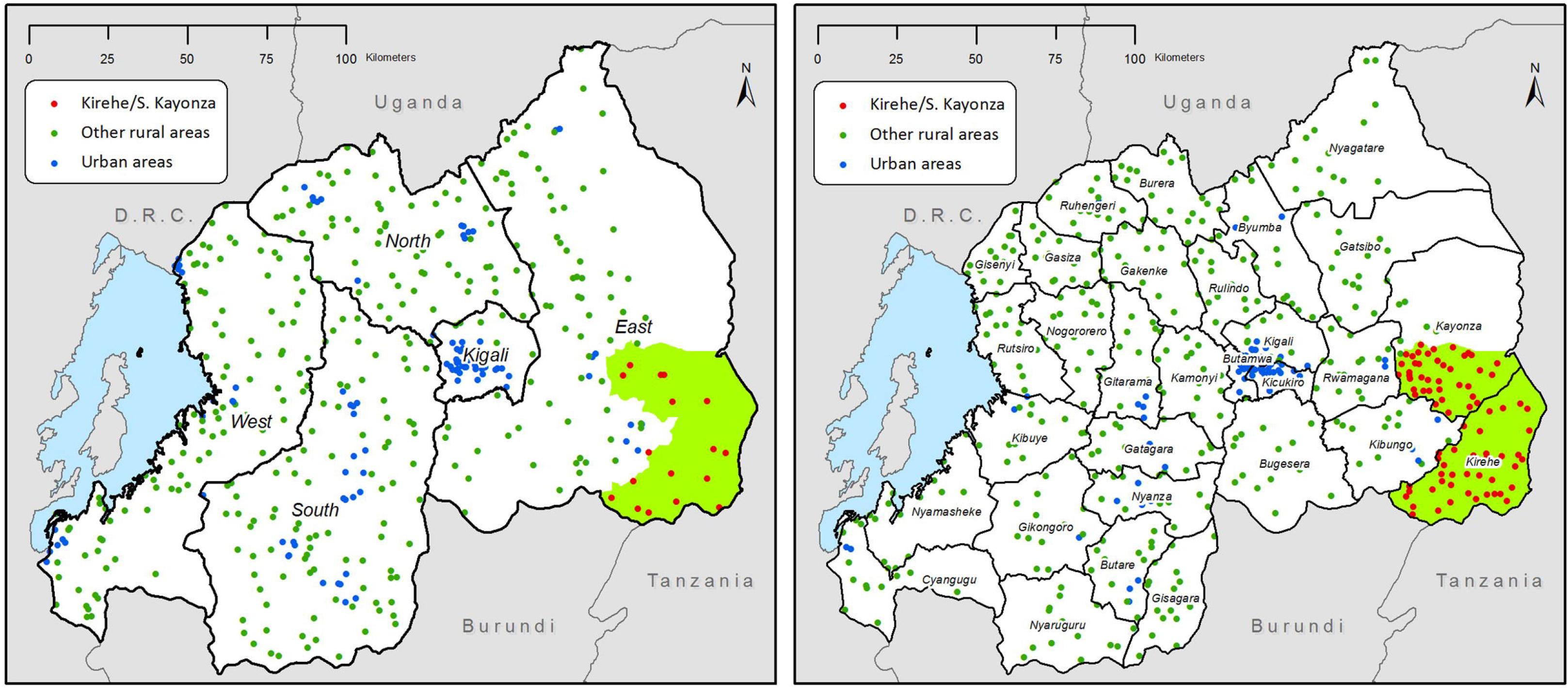
Maps of RDHS strata and primary sampling units in 2005 (left) and 2010 (right)’ and the PIH-RMOH intervention area in southeastern Rwanda (green)

### Analysis

We integrated the data from the supplemental survey with that from the 2010 RDHS as follows. Sampling probability weights were recalculated for the combined 2010 dataset. To protect respondent confidentiality, we randomly geodisplaced PSU latitude/longitude coordinates in the supplemental survey up to 5 kilometers within district boundaries according to DHS guidelines.^15^ All geographic information was linked to displaced PSU locations in a geographic information system (ArcGIS v10, ESRI). We combined latitude/longitude coordinates and rural residence information to identify respondents living in Kirehe/S. Kayzona and other rural areas. Additional PSU geographic characteristics included straight-line distance to the nearest main road in meters (downloaded from DIVA-GIS database), straight-line distance to Kigali province in kilometers (downloaded from Map Library database), elevation above sea level in meters (from RDHS), and 30-year (1971-2000) average total rainfall in millimeters during the months of January, April, July, and October (downloaded from US National Oceanic and Atmospheric Administration Climate Prediction Center). We generated comparable household wealth scores for 2010 using the principle components generated from the 2005 RDHS,^16^ and we considered a household to be “poor” if it ranked in the bottom 20% of wealth scores of the pooled 2005, 2010, and supplemental survey datasets.

We compared baseline differences in woman, household, and community (PSU) characteristics between the two comparison groups using chi-squared tests and t-tests and temporal changes in and between groups using ordinary least squares regression with a year, group, and year-by-group interaction term. We also compared the following social and geographic characteristics of sampled communities: fraction of each PSU with improved water, fraction of PSU adults who received a primary education, distance of PSU to a main road and to Kigali, elevation, and average total rainfall in specified months. Finally, we compared baseline health system outputs and population health measures at baseline.

To identify an optimal comparison group for Kirehe/S. Kayonza, we assessed PSU characteristics that might differ between the intervention and comparison areas but which would not be expected to be altered by the intervention (distance to road, distance to Kigali, elevation, and average rainfall). We generated a bias B value to capture the difference in the standard deviations between the means of the groups and an R value which is the ratio of variances in the two groups. Following Rubin, we considered the groups to be balanced if B was less than 25% and R was between 0.5 and 2.^17^ Kirehe/S. Kayonza and other rural areas were balanced by R value but not by the B bias value (B=340.9, R=0.54) (see Supplement). We performed two further analyses to try to refine our choice of comparison group. First, we limited our comparison areas to those located in Eastern Province in proximity to the intervention area (see Figure 1). Secondly, we used propensity score matching with inverse probability of treatment weights. Since neither approach identified a more appropriate comparison group (see Supplement), we compared Kirehe/S. Kayonza to all other rural areas.

We used ordinary least squares regression with group, year, and group-year interaction terms to model changes in binary health outputs and outcomes, controlling for differences in woman’s age and household wealth at baseline. We modeled change in childhood mortality rates using the DHS synthetic life-table approach which utilizes the histories of any child a mother reports to have been alive during the previous five years.^18^ Adult mortality rates were based on a five-year synthetic cohort of respondent’s sibling’s births and deaths.^19^ Expected mortality rates were calculated by standardizing mortality rates of other rural areas to the age structure in Kirehe/S. Kayonza. We estimated mean changes between 2005 and 2010 as the absolute difference in rates, and we calculated variances of trends as the sum of year-group variances. We calculated a composite coverage index (CCI) to monitor overall health care coverage across time in the intervention and comparison areas based on that proposed by Barros and Victoria (2013) but modified to exclude BCG coverage as an indicator (see Supplement).^20^ We adjusted for clustering of observations by PSU using Taylor linearized variance estimation in regression models and jackknife repeated replications to estimate variance in all other analyses.^19^ We conducted regressions in Stata v13 and mortality analyses in SAS v9.2.

### Ethics Statement

Verbal consent was obtained for all respondents before interviews took place. Protocols for the Rwanda 2005 and 2010 DHSs were approved by the Rwandan government. Protocols for the 2010 supplemental survey were reviewed and approved by the Partners HealthCare Internal Review Board (protocol #: 2009-P-001941/8) and the Rwanda National Ethics Committee.

### Role of the funding source

The funders had no role in study design, data collection, data analysis, data interpretation, writing of this report, nor the decision to submit this paper for publication. The corresponding author (DRT) had full access to all the data in the study and had final responsibility for the decision to submit for publication.

## RESULTS

### Demographic and geographical data

At baseline, women in Kirehe/S. Kayonza and other rural areas were similar in terms of age, marital status, employment, and literacy. Households in Kirehe/S. Kayonza had fewer household assets but greater access to improved water sources than in other rural areas (Table 1). Kirehe/S. Kayonza also differs from other rural areas in terms of its geography; this region has lower elevation and lower rainfall during the second half of the calendar year (Table 1). Between 2005 and 2010, households in Kirehe/S. Kayonza acquired more assets than in other rural areas and the age distribution of women shifted to the right. Access to improved water sources rose more steeply in the other rural areas during the study period (Table 1).

### Health System Outputs

After we adjusted for household wealth and woman’s age, baseline health system outputs in both populations were similar (p>0.05 for all), although vitamin A supplementation in the last six months was lower (p=0.005) and contraception use among married women was higher (p=0.011) in Kirehe/S. Kayonza (Col. D, Table 2).

**Table 2.**
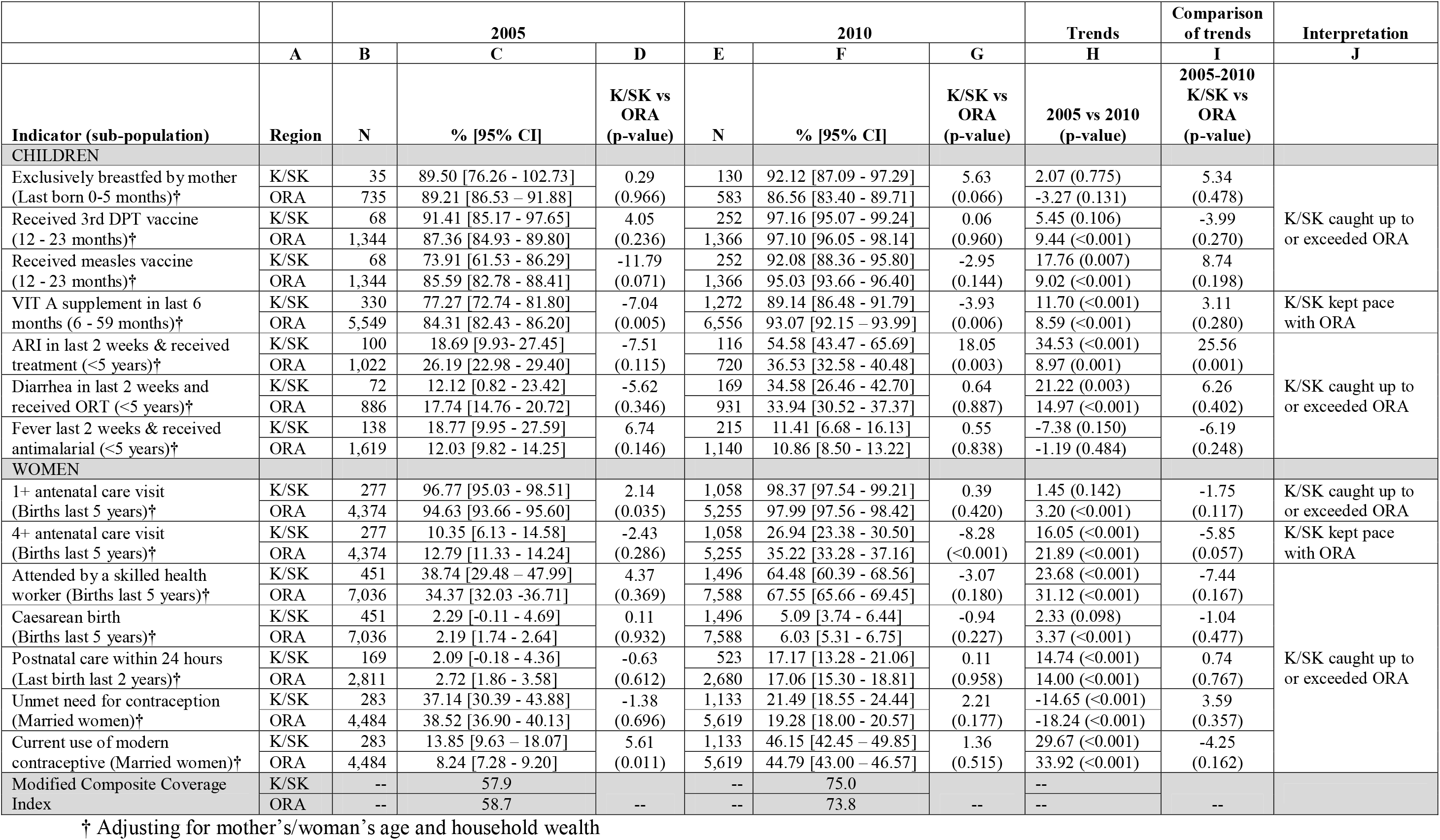
Health System Outputs 2005-2010 in Kirehe/S. Kayonza and Other Rural Areas

Most health system output indicators improved between 2005 and 2010 in both groups on most indicators (p<0.05) (Col. H, Table 2). Exceptions included exclusive breastfeeding for the first six months of life (K/SK: p=0.775, ORA: p=0.131) and treatment of fever with antimalarials (K/SK: p=0.150; ORA: p=0.484) which did not change significantly in either group over time (Col. H, Table 2). The change in the proportion of children treated for ARI (p=0.001) and in the proportion of women receiving four antenatal visits (p=0.057) was greater in Kirehe/S. Kayonza (Col. I, Table 2).

Kirehe/S. Kayonza had either caught up to (p>0.05) or exceeded (p<0.05) the performance of other rural areas in most health system outputs by 2010, with the exceptions of vitamin A supplementation (p=0.006) and the proportion of pregnant women who received four ANC visits (p<0.001) (Col. G, Table 2). Overall health system coverage as measured by the modified CCI increased from 57.9% to 75.0% in Kirehe/S. Kayonza and from 58.7% to 73.8% in other rural areas (Table 2).

### Population Health Outcomes

After adjustment, we found that most baseline measures of child health were worse in Kirehe/S. Kayonza than in other rural areas (10.99% more ARI, p=0.005; 5.41% more diarrhea, p=0.038; 11.91% more fever, p=0.003; 72 more under-five deaths per 1000 births, p=0.045), with the exception of stunting which was 5.43% more prevalent in other rural areas (p=0.018) (Col. D, Table 3). We noted improvements between 2005 and 2010 for both groups in ARI, fever, neonatal mortality, infant mortality, and under-five mortality (p<0.05 for all) (Col. H, Table 3); these improvements were greater for ARI, diarrhea, and fever in Kirehe/S. Kayonza (p<0.05 for all) (Col. J, Table 3).

Importantly, under-five mortality dropped precipitously in both groups during the study period with 147 fewer under-five deaths per 1000 live births in Kirehe/S. Kayonza (p<0.001), compared to 82 in other rural areas (p<0.001) (Col. H, Table 3). These changes represent annual reductions in under-five mortality of 12.8% and 8.9% for the intervention and other rural areas respectively (Col. I, Table 3). The greatest reductions in childhood mortality occurred among the poorest households in Kirehe/S. Kayonza with a dramatic drop in mortality from 275.4 to 89.4 deaths per 1000 live births in Kirehe/S. Kayonza (annual rate of reduction: 13.5%) compared to 152.2 to 76.2 in other rural areas (annual rate of reduction: 9.0%) (Figure 2). By 2010, there were no significant differences between Kirehe/S. Kayonza and other rural areas in terms of child health outcomes (p>0.05 for all) (Col. G, Table 3).

**Figure 2.**
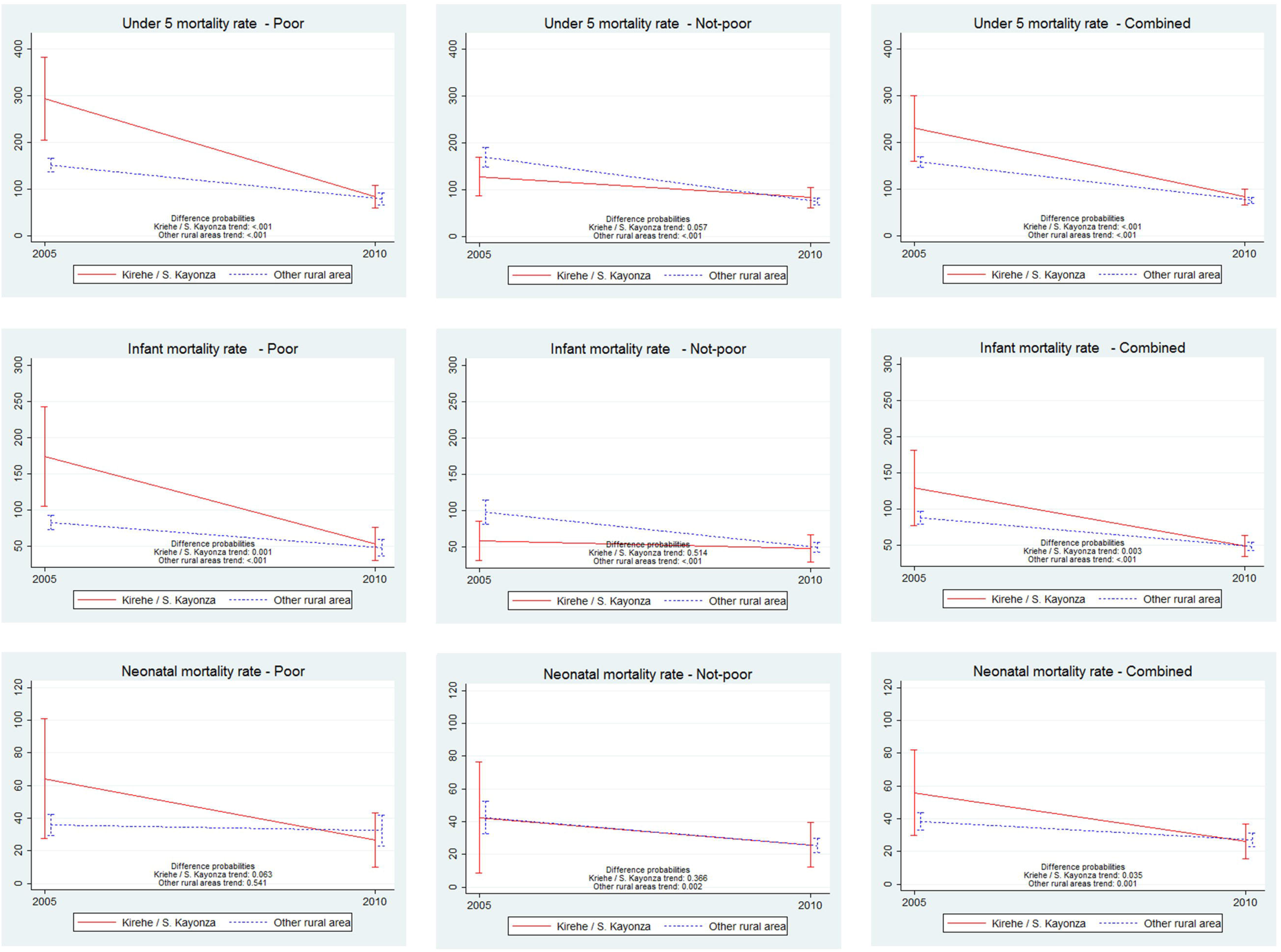
Comparison of trends in under 5’ infant’ and neonatal mortality between 2005 and 2010 in Kirehe/S. Kayonza versus Other Rural Areas’ by household wealth status

**Table 3.**
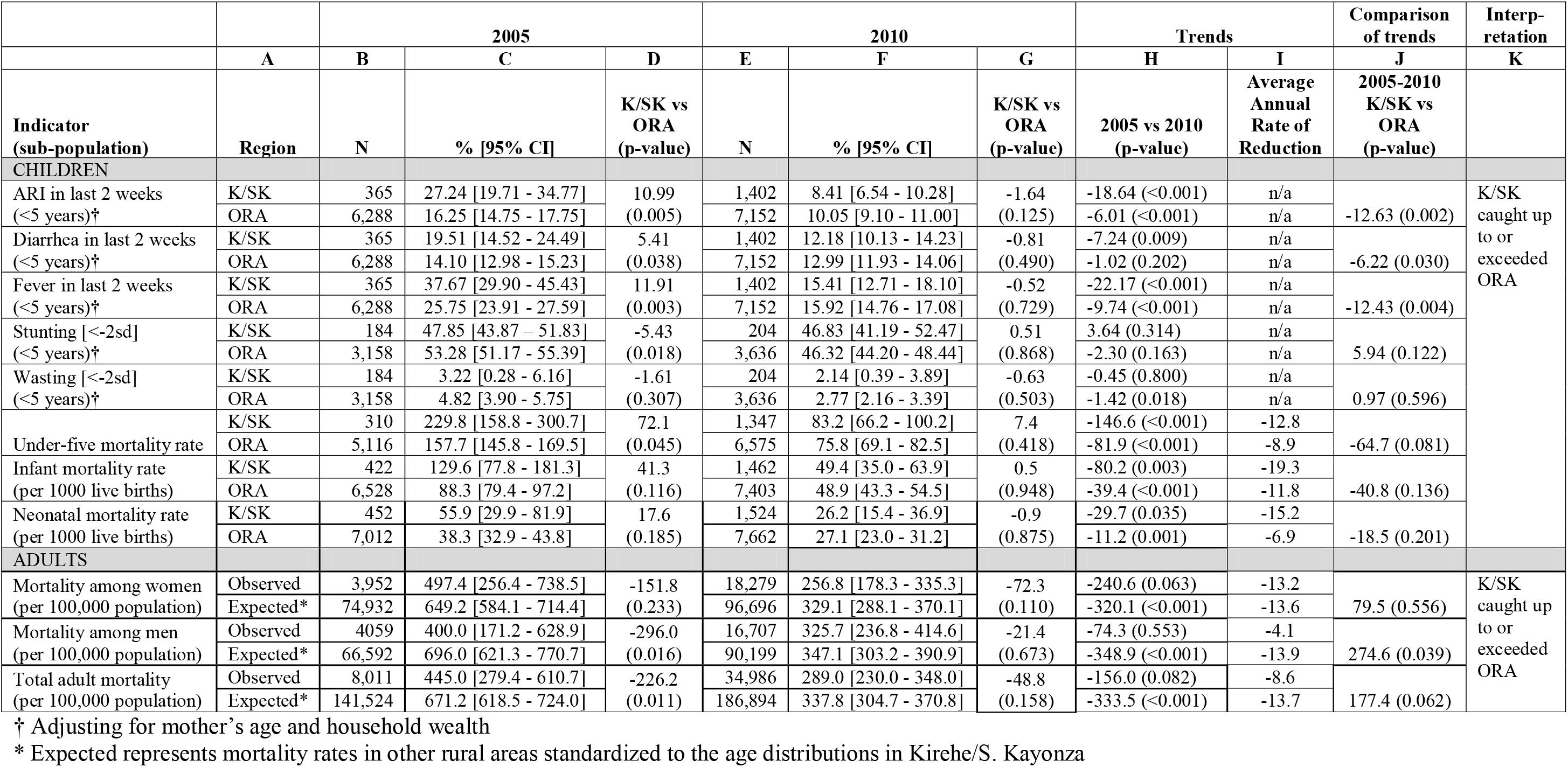
Health Outcomes 2005-2010 in Kirehe/S. Kayonza and Other Rural Areas’ adjusting for mother’s age and household wealth

In 2005, the standardized male adult mortality rate was lower in Kirehe/S. Kayonza than in other rural areas (p=0.016) and there was no difference in rates for women (p=0.233) (Col. D, Table 3). Both female and male adult mortality in Kirehe/S. Kayonza dropped between 2005 and 2010 (from 497 to 257 deaths per 1000 among women, and from 400 to 326 deaths per 1000 among men) (Col. C and F, Table 3), and there was no difference in the change in standardized adult mortality rates between the two groups (p=0.062) (Col. J, Table 3).

## DISCUSSION

Between 2005 and 2010, coverage of health system outputs improved dramatically in both the area of southeast Rwanda targeted by the intervention described here and in other rural areas of the country. This increased coverage was accompanied by steep declines in adult, under-five, infant, and neonatal mortality in both settings. Most of these gains were more extreme in the intervention area; the annual rate of reduction in under-five mortality, for example, was over 12% in the Kirehe/Kayonza compared to 8.9% in other rural areas. The difference in this and other rates of decline in population outcomes did not meet statistical significance, however, in part because the 2005 RDHS was underpowered to detect subregional differences and we did not oversample the intervention area at baseline.

These data are consistent with the findings from a recent overview of global trends in child mortality which identified Rwanda as a top performer worldwide in reducing under-five mortality between 2000 and 2015.^21^ Though Rwanda is classified as a least developed country, its national 9.9% annual rate of reduction in under-five deaths is surpassed only by the middle-high income nation of the Maldives.^22^ This reduction is more than twice the global rate of 4.4% during the same time period.^22^

Multiple studies in other settings suggest that national health gains can often mask substantial heterogeneity in health system performance and outcomes.^23,24^ In many settings, impoverished and/or geographically inaccessible areas have experienced slower progress in achieving health goals.^25,26^ In other studies, socioeconomic status did not correlate well with health system performance.^27^ Our results show that the historically unprecedented decline in health indicators extends not only to urban and wealthier areas of the country but is also possible among its poorest and most geographically isolated residents. Notably, the decline in under-five mortality that we observed was steepest among the lowest two wealth quintiles in the Kirehe/S. Kayonza region where the intervention included specific components (subsidies of insurance premiums and co-pays, nutrition support, and compensated village based community health workers) designed to address inequities in access to care.

How were the remarkable improvements in health outcomes made in the regions and socioeconomic groups we assessed? Several previous reports describe the major components of Rwanda’s national health strategy during this period of success.^21,28^ Like Rwanda’s plan to improve health outcomes throughout the country, our intervention was deliberately comprehensive and it is challenging to disentangle specific components that contributed to the health gains observed. Interestingly, despite the fact that under-five mortality was almost one third lower in other rural areas than it was in Kirehe/S. Kayonza in 2005, the modified composite coverage index of the set of interventions thought to reduce under-five mortality was very similar between the groups. Nonetheless, Kirehe/S. Kayonza initially trailed the other rural areas in two interventions most likely to target common causes of death among children in lower- and middle-income countries (LMICs): case management of pneumonia and diarrhea. The CCI increased similarly in both groups, but Kirehe/S. Kayonza had closed the gap in case management for diarrhea and achieved higher coverage for exclusive breast feeding and case management of ARI by 2010. In contrast, other rural areas outperformed Kirehe/S. Kayonza in measles vaccination, vitamin A supplementation, ANC4 and skilled birth attendants, all of which might be expected to have a less direct impact on overall mortality.

In addition to a focus on equity, the construction or renovation of two district hospitals, and higher coverage of horizontal interventions addressing major causes of child death. Specifically, the Kirehe/S. Kayonza intervention combined a rigorous program of compensated village-based community health workers with improvements in infrastructure and staffing of the facilities to which CHWs referred patients for care. The currently accepted indicators of coverage assess the frequency, but not the quality of the care provided, are therefore coarse tools by which to measure the impact of the range of interventions embodied in the six WHO building blocks. For example, treatment of malaria with an antipyretic rather than artemesin meets the criteria for case management of fever as assessed in the DHS but is unlikely to have a major impact on child mortality. Recognizing the challenges of designing an evaluation that captures the relative effects of these components of our care model, we suggest that the impact of the Kirehe/S. Kayonza intervention is partly attributable to the coordinated strengthening of services at multiple levels (community, health centers, and hospitals) with a focus on quality of care.^29^

We note several important limitations to this study. First, mother-reports may be imperfect measures of illness and treatment, especially for such indicators as acute respiratory infections including pneumonia and whether the child received antibiotic treatment.^30^ Second, the sample size at baseline was small. Dwyer-Lindgren and colleagues have shown that under-five mortality estimates derived from birth histories with sample sizes under 500 can be biased, and usually underestimate the true value of mortality.^31^ They also show that underestimation of mortality tends to occur when estimates are based on surveys conducted in high mortality settings. With a sample size of only 359 birth records in the 2005 survey of the very high mortality intervention region, it is possible that the true Kirehe/S. Kayonza baseline under-five mortality rate was even higher and the decline between 2005 and 2010 steeper than we estimated.

Third, the cross-sectional nature of the surveys did not allow for tracking changes in individuals, households, or communities. Anecdotal evidence suggests that a number of chronically sick individuals and their families from nearby districts moved to Kirehe/S. Kayonza after health system improvements began. The opening of new health facilities and establishment of the PIH-Rwanda headquarters in Kirehe/S. Kayonza also attracted new, better-educated employees to the region and stimulated related commerce. The cross-sectional study design does not allow for determining to what extent changes in health were driven by the health system strengthening intervention in the baseline population, compared to other demographic changes in the population that occurred during the intervention period. Fourth, we were only able to measure WHO building block indicators available in the RDHS questionnaire; data about perception or quality of care, for example, could not be included. As in any household questionnaire, selection and recall biases may have also affected estimates. Fifth, we could not identify an ideal comparison group for Kirehe/S. Kayonza at baseline to be able to conduct a strict difference-in-differences analysis of the effect of the RMOH-PIH intervention because the criteria for similar groups at baseline was not met.

## CONCLUSION

Rwanda experienced historic health improvements between 2005 and 2010, and those improvements were even more pronounced in Kirehe/S. Kayonza where RMOH-PIH rolled out an integrated health system strengthening intervention in 2005. This area had substantially poorer performance on population health outcomes in 2005, but was able to catch up to or exceed a fast moving target on 23 of 25 indicators at the end of the first five years of the RMOH-PIH intervention. Furthermore, the drop in under-five mortality in Kirehe/S. Kayonza was higher than that of the historic drop experienced nationally and in other rural areas. Although we are not able to attribute health improvements in Kirehe/S. Kayonza to the RMOH-PIH program alone, the RMOH-PIH program likely played a key role in these monumental health achievements.

## Acknowledgements

We thank ICF International and the Rwanda Ministry of Health for collecting these data and supporting the cleaning of supplemental survey data in Kirehe and Southern Kayonza districts. We also thank Paulin Basinga, Jeanine Condo, Saleh Niyonzima, Adolphe Karamaga, Cathy Mugeni, and Fidele Ngabo who contributed to the design and implementation of these methods, and James Robins and Bethany Hedt-Gauthier for discussions about statistical modeling. Finally, deepest thanks the hundreds of staff and health care providers who have worked in Kirehe and Southern Kayonza over the last decade to deliver quality care and continually pursue improvements in Rwanda’s health system. This paper is a product of a collaborative working group that was led by Harvard Medical School. The content of this article is solely the responsibility of the authors and does not necessarily represent the official views of the institutions to which the authors are affiliated. Data collection for this project was paid for by the Doris Duke Charitable Foundation through the Population Health Implementation & Training (PHIT) Partnerships that link implementation research and training directly to health care delivery.

